# Inhibitors of anti-apoptotic Bcl-2 family proteins exhibit potent and broad-spectrum anti-mammarenavirus activity via cell cycle arrest at G0/G1 phase

**DOI:** 10.1101/2021.08.16.456587

**Authors:** Yu-Jin Kim, Haydar Witwit, Beatrice Cubitt, Juan C. de la Torre

**Author notes:** Corresponding author: Department of Immunology and Microbiology, The Scripps Research Institute, La Jolla, CA 92037.

## Abstract

Targeting host factors is a promising strategy to develop broad-spectrum antiviral drugs. Drugs targeting anti-apoptotic Bcl-2 family proteins that were originally developed as tumor suppressors have been reported to inhibit multiplication of different types of viruses. However, the mechanisms whereby Bcl-2 inhibitors exert their antiviral activity remain poorly understood. In this study, we have investigated the mechanisms by which obatoclax (OLX) and ABT-737 Bcl-2 inhibitors exhibited a potent antiviral activity against the mammarenavirus lymphocytic choriomeningitis virus (LCMV). OLX and ABT-737 potent anti-LCMV activity was not associated with their pro-apoptotic properties, but rather their ability of inducing cell arrest at G0/G1 phase. OLX and ABT-737 mediated inhibition of Bcl-2 correlated with reduced expression levels of thymidine kinase 1 (TK1), cyclin A2 (CCNA2), and cyclin B1 (CCNB1) cell cycle regulators. In addition, siRNA-mediated knock down of TK1, CCNA2, and CCNB1 resulted in reduced levels of LCMV multiplication. The antiviral activity exerted by Bcl-2 inhibitors correlated with reduced levels of viral RNA synthesis at early times of infection. Importantly, ABT-737 exhibited moderate efficacy in a mouse model of LCMV infection, and Bcl-2 inhibitors displayed broad-spectrum antiviral activities against different mammarenaviruses and SARS-CoV-2. Our results suggest that Bcl-2 inhibitors, actively being explored as anti-cancer therapeutics, might be repositioned as broad-spectrum antivirals.

**IMPORTANCE:** Anti-apoptotic Bcl-2 inhibitors have been shown to exert potent antiviral activities against various types of viruses via mechanisms that are currently poorly understood. This study has revealed that Bcl-2 inhibitors mediated cell cycle arrest at the G0/G1 phase, rather than their pro-apoptotic activity, plays a critical role in blocking mammarenavirus multiplication in cultured cells. In addition, we show that Bcl-2 inhibitor ABT-737 exhibited moderate anti-mammarenavirus activity *in vivo*, and that Bcl-2 inhibitors displayed broad-spectrum antiviral activities against different mammarenaviruses and SARS-CoV-2. Our results suggest that Bcl-2 inhibitors, actively being explored as anti-cancer therapeutics, might be repositioned as broad-spectrum antivirals.

## INTRODUCTION

The Bcl-2 family proteins are critical regulators of apoptosis and are classified into three groups: anti-apoptotic proteins (Bcl-2, -X_L_, -W, MCL-1, BFL-1/A1), pro-apoptotic pore-forming proteins (BAX, BAK, BOK), and pro-apoptotic BH3-only proteins (BAD, BID, BIK, BIM, BMF, HRK, NOXA, PUMA) (1). Upon apoptotic stimuli, pro-apoptotic pore-forming members generate pores within the mitochondrial outer membrane, thus promoting mitochondrial outer membrane permeabilization (MOMP). MOMP results in the release of mitochondrial intermembrane space (MIS) proteins including cytochrome c and SMAC/DIABLO into the cytoplasm, leading to the activation of the caspase cascade and apoptotic cell death (2). The anti-apoptotic group of Bcl-2 proteins inhibit both pro-apoptotic groups, pore-forming and BH3-only members, by direct binding to BH3 domains and blocking the progression of the apoptotic signaling cascade before activation of the effector caspases takes place (1, 3). Due to their role in cell survival, anti-apoptotic Bcl-2 proteins are considered as attractive targets of tumor suppressing agents and a number of inhibitors of Bcl-2 proteins have been developed as chemotherapeutic candidates for cancer treatment (4). Anti-apoptotic Bcl-2 inhibitors mimic the BH3 domain of the pro-apoptotic Bcl-2 members and occupy the BH3-binding site in anti-apoptotic Bcl-2 proteins, promoting activities of pro-apoptotic Bcl-2 proteins and apoptosis progression (5). Besides promoting apoptosis, anti-apoptotic Bcl-2 inhibitors have been reported to exert antiviral activities against various types of viruses (6-12). However, the mechanisms by which Bcl-2 inhibitors exert their antiviral activities are not well understood. Several Bcl-2 inhibitors can arrest cells at the G0/G1 phase of the cell cycle, which disrupts normal proliferation and migration of tumor cells (13-15). Notably, several viruses encode proteins that interact with cell cycle regulating proteins and alter cell cycle progression in different ways (16), but how these interactions can impact virus multiplication remains to be elucidated.

In a previous drug screening, we identified OLX, an antagonist of anti-apoptotic Bcl-2 proteins, as a potent anti-mammarenaviral drug (17). To investigate the mechanisms whereby Bcl-2 inhibitors could exert their anti-mammarenavirus activity, we selected the Bcl-2 inhibitors OLX and ABT-737 due to their potent inhibitory effect on the activity of anti-apoptotic Bcl-2 proteins *in vitro* (18, 19) and *in vivo* (20-24). We found that the antiviral activity exerted by OLX and ABT-737 against the prototypic mammarenavirus lymphocytic choriomeningitis virus (LCMV) was independent of apoptosis and correlated with OLX and ABT-737 induced cell cycle arrest at the G0/G1 phase, which was associated with reduced expression levels of multiple cell cycle regulators including thymidine kinase 1 (TK1), cyclin A2 (CCNA2), and cyclin B1 (CCNB1). Notably, the altered cellular environment caused by cell cycle arrest at G0/G1 interfered with LCMV RNA synthesis at early times of infection. Importantly, ABT-737 exhibited modest in vivo antiviral efficacy reflected by mice infected with strain clone 13 of LCMV and treated with ABT-737 exhibiting less weight loss and lower viral load compared to vehicle treated infected mice. Both OLX and ABT-737 exhibited also a potent antiviral effect against the mammarenaviruses Junin (JUNV), the causative agent of Argentine hemorrhagic fever (AHF), and Lassa virus live-attenuated vaccine candidate reassortant ML29, as well as against SARS-CoV-2, suggesting a broad-spectrum antiviral activity of Bcl-2 inhibitors. Our findings have uncovered a novel mechanism by which Bcl-2 inhibitors can exert their antiviral activity, highlighting modulation of the cell cycle as a potential target for development of broad-spectrum antiviral therapeutics.

## RESULTS

### Effect of Bcl-2 inhibitors on LCMV multiplication in cultured cells

To investigate whether OLX-mediated anti-mammarenavirus activity was also observed with other inhibitors of anti-apoptotic Bcl-2 family proteins, we compared the effect of the Bcl-2 inhibitors OLX and ABT-737 on multiplication of a recombinant LCMV expressing the ZsGreen (ZsG) reporter gene (rLCMV/ZsG) in A549 cells. Cells were treated with 3-fold serial dilutions of each Bcl-2 inhibitor starting at 2 hours before infection. At 48 hours post-infection, virus multiplication was assessed based on ZsG expression levels that were normalized to vehicle (DMSO) treated controls (Fig. 1A). We also examined the dose-dependent effect of OLX and ABT-737 on cell viability (Fig. 1B). Both OLX and ABT-737 exhibited a potent dose-dependent inhibitory effect on LCMV multiplication with half-maximal effective concentrations (EC_50_) of 0.01 µM and 0.40 µM, respectively, and selectivity index (SI; SI = CC_50_/EC_50_) values of 113.60 (OLX) and 49.41 (ABT-737) (Fig. 1C).

**Figure 1.**
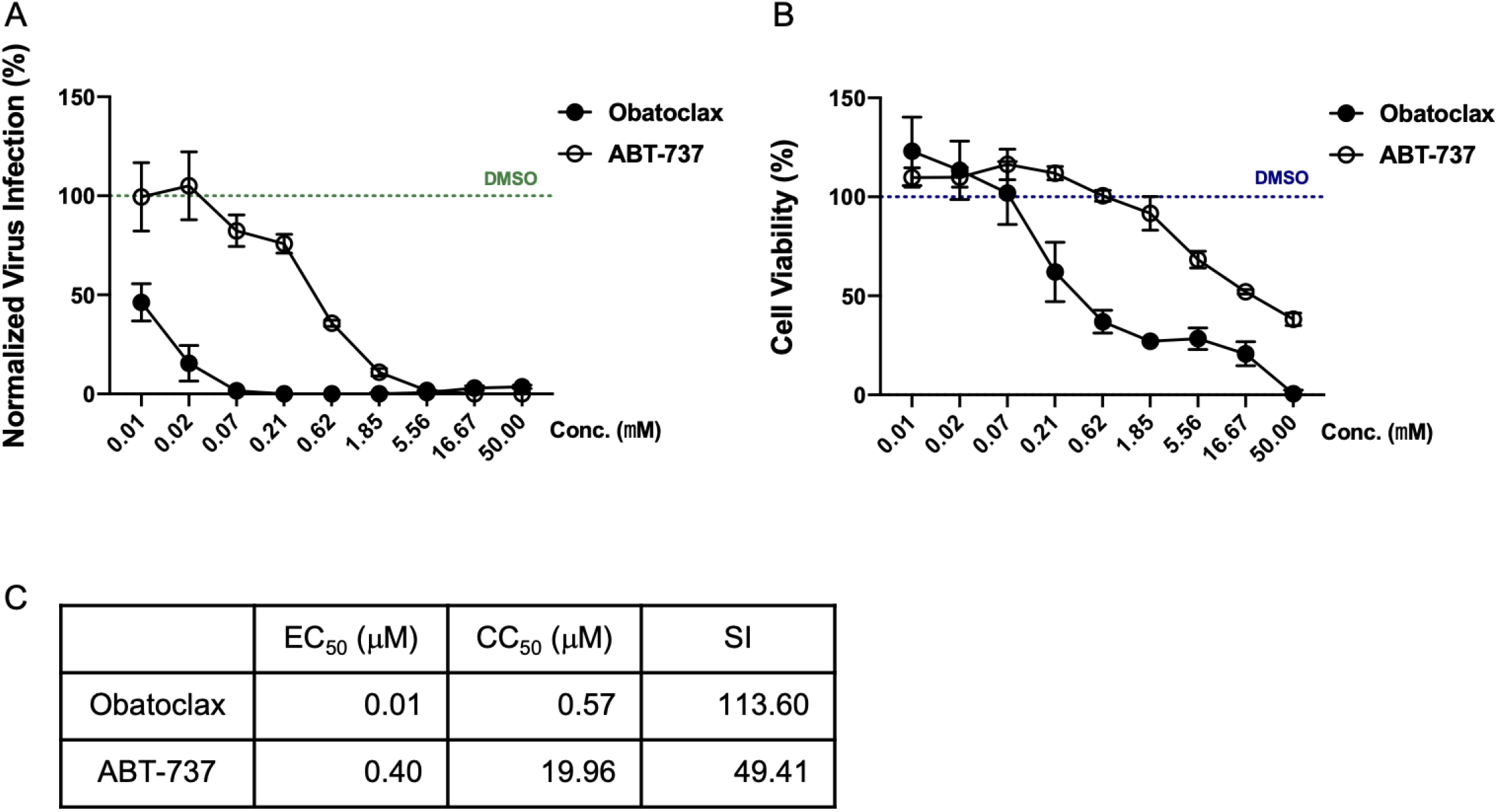
Antiviral activity of apoptosis inducing Bcl-2 inhibitors. (A) Virus propagation in the presence of OLX or ABT-737. A549 cells were treated with serial dilutions of each compound starting 2 hours prior to infection (MOI = 0.01) with rLCMV/ZsG. At 48 hpi ZsG expression levels were measured and normalized to vehicle control (DMSO) treated controls. (B) Cell viability was determined using the CellTiter-Glo assay at 48 hours compound treatment. Results were normalized to vehicle (DMSO) treated controls. (C) EC_50_, CC_50_, and SI (CC_50_/EC_50_) were calculated for OLX and ABT-737.

OLX and ABT-737 were developed as apoptosis inducing agents, via their inhibitory activities on Bcl-2 family proteins (18-24), to promote cell death of cancer cells. We therefore investigated whether apoptosis induction caused by Bcl-2 inhibition contributed to the antiviral activities of OLX and ABT-737. We first confirmed that ABT-737 treatment of A549 cells resulted in an increase of both early (Annexin V^+^ / 7-AAD^-^) and late (Annexin V^+^ / 7-AAD^+^) apoptotic cells (Fig. 2A). Under our experimental conditions, OLX exhibited broad-spectrum autofluorescence in a dose-dependent manner, which prevented us to assess its effect on cell apoptosis by flow cytometry analysis. We further confirmed the apoptosis inducing effects of both Bcl-2 inhibitors by detection of caspase activity. OLX and ABT-737 significantly increased the activity of caspase-3, a critical component of the apoptotic pathway (Fig. 2B). However, treatment with either the pan-caspase inhibitor zVAD, or with the caspase-3 specific inhibitor zDEVD did not affect the antiviral activity of OLX or ABT-737 (Fig. 2C). To rule out the possibility that LCMV infection could interfere with the apoptotic pathway, we determined levels of caspase activity in LCMV-infected cells treated with OLX or ABT-737, or vehicle (DMSO) control, in the presence and absence of the pan-caspase inhibitor zVAD. LCMV infection did not affect Bcl-2 inhibitor-mediated caspase activation, and zVAD effectively inhibited caspase activity in LCMV-infected cells (Fig. 2D).

**Figure 2.**
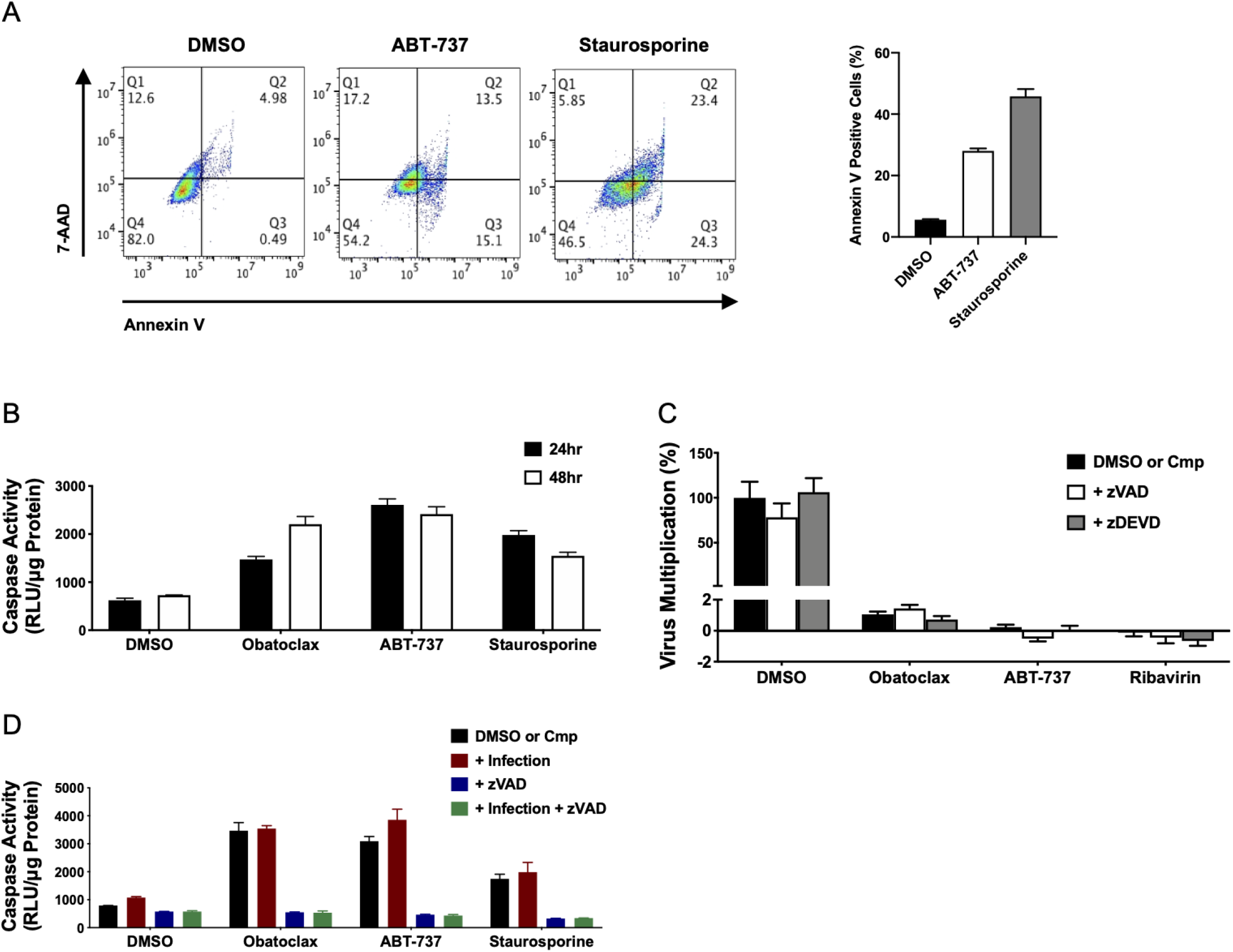
Effects of Bcl-2 inhibitor-induced apoptosis on LCMV multiplication. (A) A549 cells were treated with the indicated compounds and concentrations for 48 hours. Harvested cells were incubated with Annexin V conjugate (Alexa Fluor 488) and 7-AAD and analyzed by flow cytometry. Plots are the representative gatings of each compound (left panel). The graph reveals Annexin V positive cell populations (Q2 + Q3) (right panel). (B) Whole cell lysates were collected at the indicated times and caspase-3/7 activity was determined and normalized to total protein amount in the cell lysate. (C) A549 cells were treated with the indicated compounds 2 hours before infection with rLCMV/ZsG. At 48 hpi, ZsG expression levels were determined and normalized to vehicle treated control. (D) A549 cells were treated with the compounds for 2 hours followed by infection with rLCMV/ZsG. At 48 hpi, caspase-3/7 activity was determined in whole cell lysates and values were normalized to total protein in cell lysates. Treatment concentration: OLX (0 .1 µM), ABT-737 (2 µM), staurosporine (1 µM), ribavirin (100 µM), zVAD (100 µM), and zDEVD (100 µM).

### Cell cycle arrest at G0/G1 phase correlated with the anti-LCMV activity mediated by OLX and ABT-737 Bcl-2 inhibitors

The mechanisms by which Bcl-2 targeting compounds exert their physiological effects are not well understood and likely involve different cellular responses including cell cycle progression (13-15, 25), autophagy (26), cell metabolism (27), cell migration (28) and apoptosis (29). Bcl-2 inhibitors have been reported to cause cell cycle arrest at G0/G1 phase (13-15). We therefore tested whether altered regulation of the cell cycle contributed to Bcl-2 inhibitor-mediated antiviral activity. We analyzed cell cycle progression in the presence of the indicated drugs at 48 hours post treatment (Fig. 3A). Treatment with ABT-737 resulted in substantial cell arrest at the G0/G1 phase. As a control, we used the CDK inhibitor dinaciclib, which showed the expected effect in cell cycle progression, increasing the G2/M phase population. Due to technical difficulties in flow cytometry analysis caused by OLX autofluorescence property, we examined the impact of OLX on cell cycle modulation by determining its effects on protein expression levels of cell cycle regulators (Fig. 3B). OLX and ABT-737 exhibited similar effects on the expression pattern of cell cycle regulators, reducing expression levels of thymidine kinase 1 (TK1), cyclin A2 (CCNA2), and cyclin B1 (CCNB1). The G2/M arresting reagent, dinaciclib, resulted in significant increase of TK1 and moderated reduction of CCNA2 and CCNB1. We included treatment with the nucleoside analog ribavirin as a reference for the effects of a broad-spectrum antiviral (30). Ribavirin treatment caused only minor reduction of CCNB1 expression. We next tested whether these cell cycle regulators contributed to LCMV multiplication. siRNA-mediated knock-down of TK1, CCNA2, and CCNB1, as well as Bcl-2, resulted in significant reduction of virus multiplication (Fig. 3C). Knock-down level of each factor was confirmed by western blot analysis, revealing that Bcl-2 knock-down caused significant decreased in expression levels of TK1, CCNA2, and CCNB1 (Fig. 3D), suggesting that Bcl-2 is an upstream modulator of these cell cycle regulators associated with progression from S to G2/M phase. Modulation of the cell cycle has been documented as a strategy used by several viruses to enhance their replication (16). To assess whether LCMV infection influenced cell cycle regulation, we infected cells with LCMV in the presence of the indicated compounds and at 48 hours post-infection (hpi) determined the distribution of cells at each phase of the cell cycle (Fig. 3E). LCMV infection did not affect cell cycle regulation of untreated or ABT-737 treated cells.

**Figure 3.**
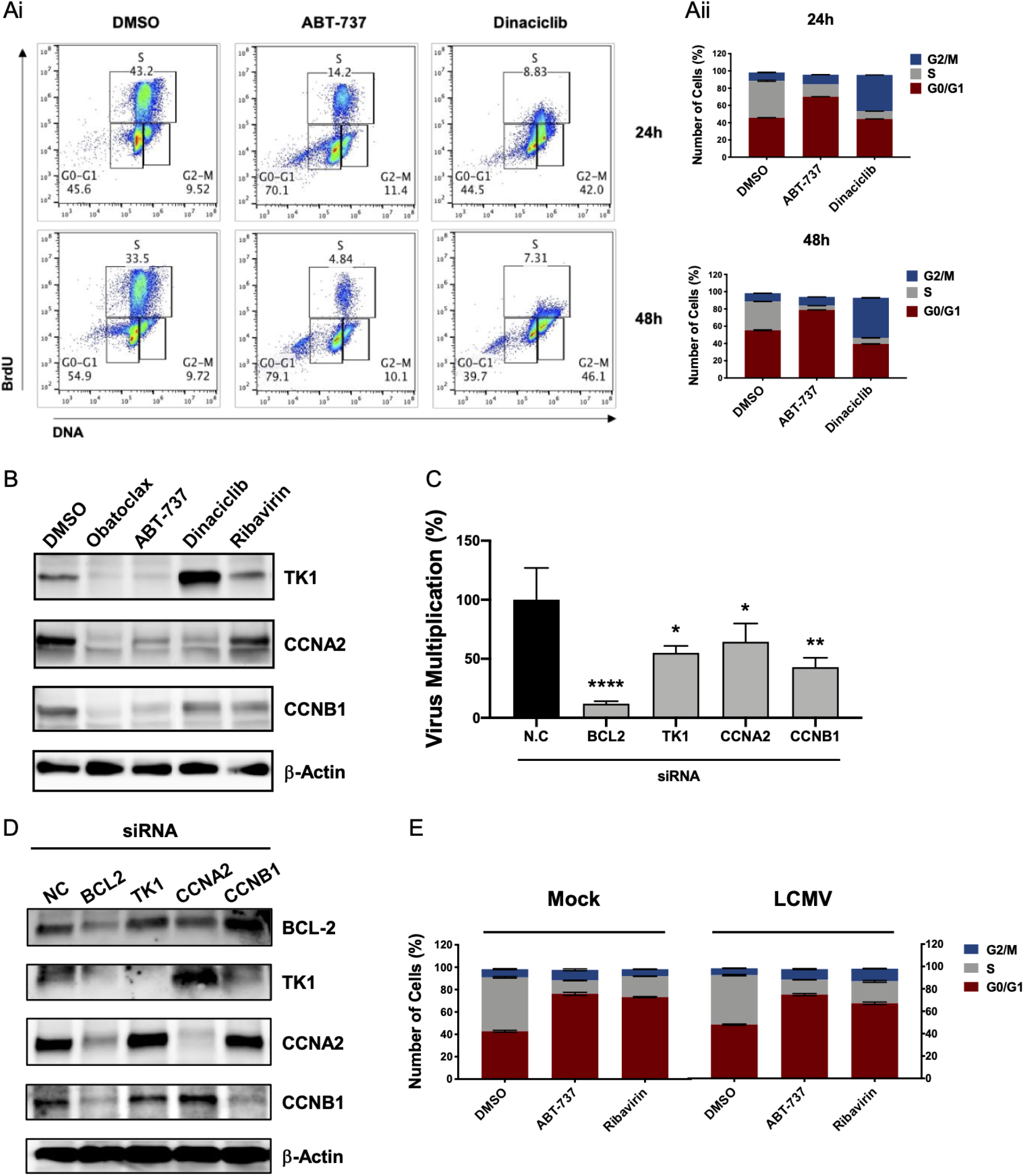
Effects of Bcl-2 inhibitor-induced G0/G1 arrest on LCMV multiplication. (A) A549 cells were treated with the indicated compounds and concentrations for the indicated times, followed by addition of BrdU to the cell culture medium for 1 hour before harvesting cells. In fixed cells, the cell cycle positions and active DNA synthesis were analyzed by the correlated levels of total DNA and incorporated BrdU using flow cytometry. Ai. Plots are representative gatings for each compound treatment (left panel). Aii. Cells (%) at each phase of the cell cycle at 24 and 48 h treatment with each compound. (B) Whole cell lysates were collected at 48 hours post-treatment and protein expression levels were determined by western blotting. (C-D) 293T cells were transfected with the indicated siRNAs and for 4 hours later infected (MOI = 0.01) with rLCMV/ZsG. At 48 hpi, levels of infectious progeny in tissue culture supernatants (C) and protein expression levels (D) were determined by FFA and western blotting, respectively. (E) A549 cells were pre-treated with the indicated compounds and infected with rLCMV/ZsG. At 48 hpi, cell cycle was analyzed using the BrdU assay. Treatments: OLX (0.1 µM), ABT-737 (2 µM), dinaciclib (5 µM), ribavirin (100 µM). Statistical significance was calculated by analysis of variance (ANOVA) (* *p* < 0.05, ** *p* < 0.002, *** *p* < 0.0002, and **** *p* < 0.00001).

### Effect of Bcl-2 inhibition on LCMV RNA synthesis

We previously reported that OLX treatment caused a significant reduction on LCMV and LASV RNA levels at 48 hpi (17). This finding, however, reflected the effect of OLX on the steady state levels of viral RNA over multiple rounds of infection. To gain a better understanding on how Bcl-2 inhibitors might modulate viral RNA synthesis, we examined the effect of OLX and ABT-737 on viral RNA synthesis at early times of infection. We treated cells with the indicated compounds for 24 hours before LCMV infection and determined levels of viral genome and anti-genome RNA species at 2 and 4 hpi (Fig. 4). Bcl-2 inhibitors did not affect significantly viral RNA synthesis at 2 hpi. In contrast, at 4 hpi, levels of both genomic and anti-genomic viral RNA were significantly reduced in OLX and ABT-737 treated cells (Fig. 4A and B). Ribavirin slightly increased levels of genomic and anti-genomic viral RNA at 2 hpi, but at 4 hpi exhibited also an inhibitory effect in viral RNA synthesis.

**Figure 4.**
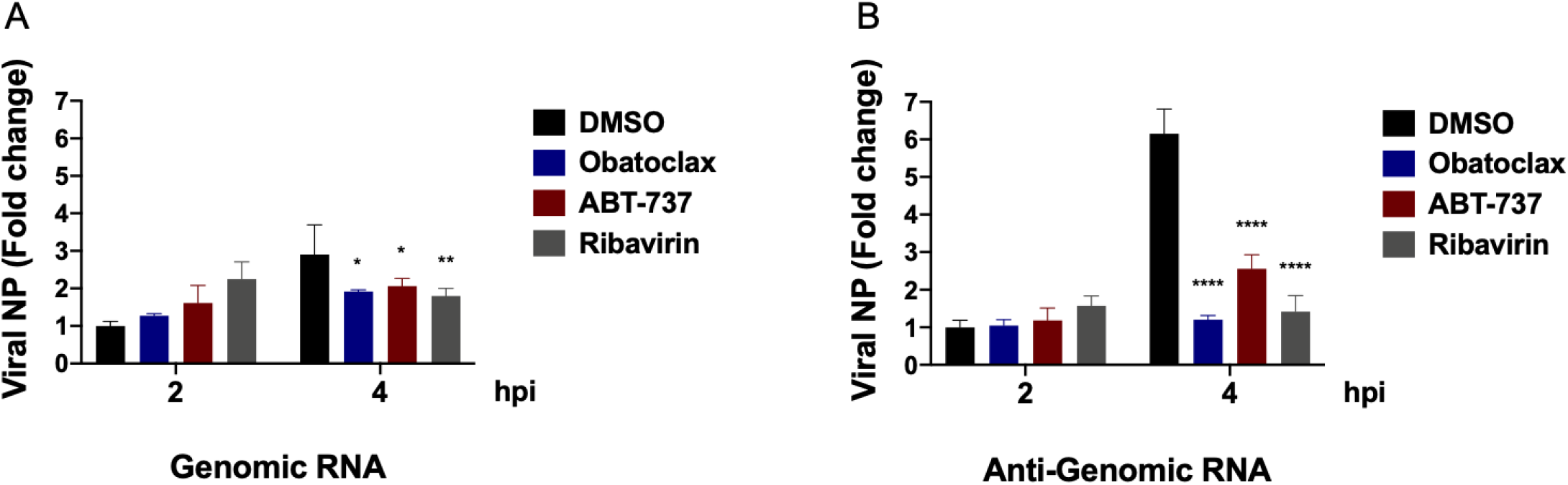
Effects of Bcl-2 inhibitors on LCMV RNA synthesis at early times of infection. (A-B) After 24 hours pre-treatment of the compounds, A549 cells were infected with rLCMV/ZsG (MOI = 3) and then the compounds were re-added. At the indicated time points, total cellular RNA was purified. Equal amounts of RNA (1 µg) were used to generate complementary DNAs (cDNAs), using strand-specific primers. Genomic (A) and anti-genomic strand (B) of cDNAs were used in quantitative polymerase chain reaction (qPCR) to determine the levels of viral nucleoprotein (NP) gene. Treatment: OLX (0.1 µM), ABT-737 (2 µM), ribavirin (100 µM). Statistical significance was calculated by analysis of variance (ANOVA) (* *p* < 0.05, ** *p* < 0.002, *** *p* < 0.0002, and **** *p* < 0.00001).

### Effect of the Bcl-2 inhibitor ABT-737 on LCMV infection *in vivo*

To assess the antiviral activity of Bcl-2 inhibitors *in vivo*, we used a mouse model of LCMV infection. We infected adult C57BL/6 mice with a high dose (2 × 10^6^ pfu/mouse) of the immunosuppressive CL-13 variant of the Armstrong strain of LCMV. We selected ABT-737 for this experiment due to its better characterized toxicology and pharmacology. Infection of adult C57BL/6 mice with a high dose of CL-13 causes transient body weight loss and persistent viremia (31). ABT-737 treated mice showed less body weight loss and faster recovery compared to the vehicle control group (Fig. 5A). Moreover, we observed moderate reduction of viral load in ABT-737 treated mice (Fig. 5B).

**Figure 5.**
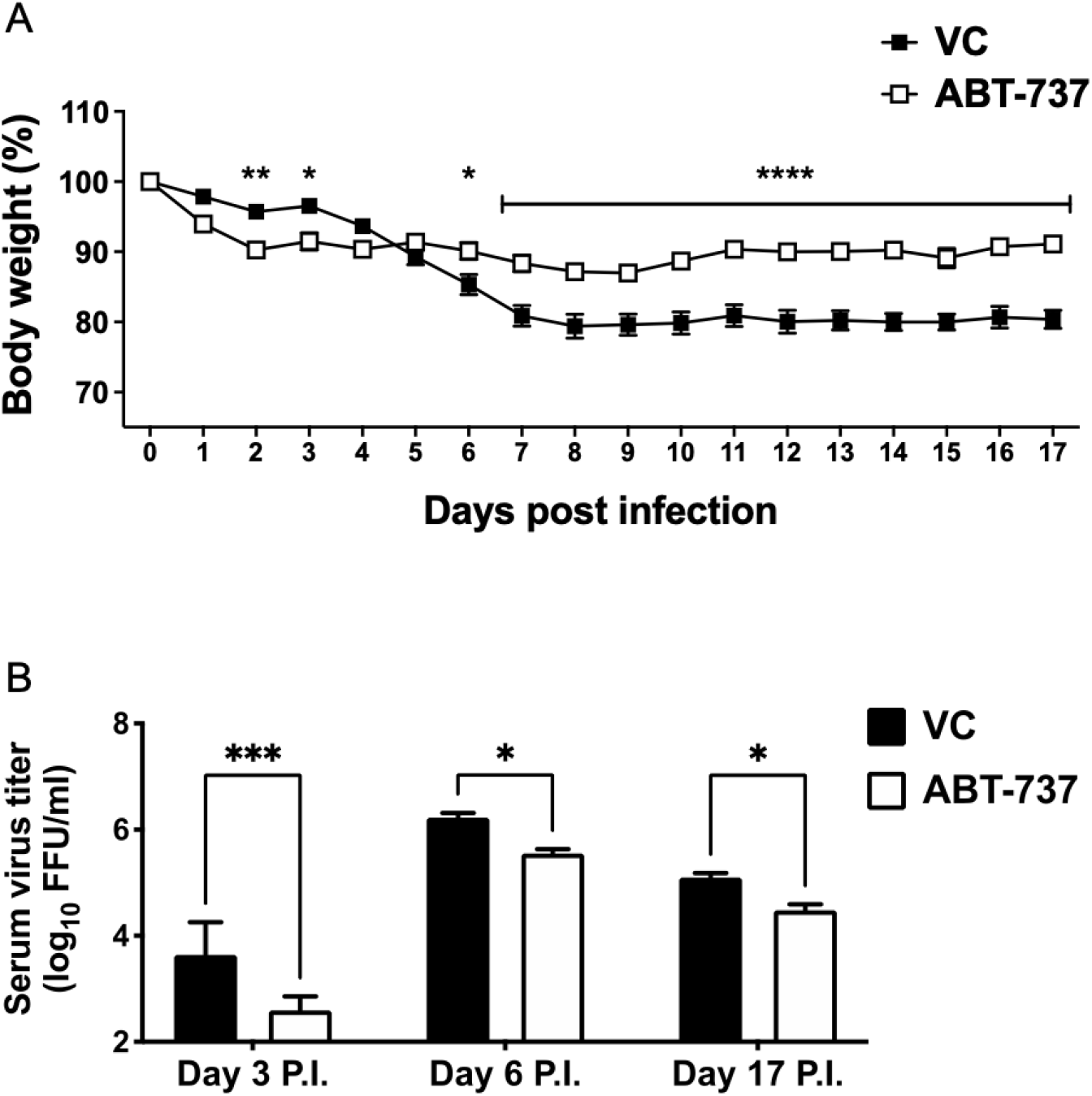
Antiviral effects of the Bcl-2 inhibitor in vivo. (A-B) Adult C58BL/6 mice (n = 4/group) were treated with ABT-737 (20mg/kg/day; intraperitoneal route) or vehicle for 17 days. At day 0, mice were infected with rCl-13 (2 × 10^6^ pfu/mouse). Body weight changes (A) and serum virus titers (B) were determined at the indicated time points. Statistical significance was calculated by analysis of variance (ANOVA) (* *p* < 0.05, ** *p* < 0.002, *** *p* < 0.0002, and **** *p* < 0.00001).

### Assessing the broad-spectrum antiviral activity of Bcl-2 inhibitors

Host targeting antiviral drugs are likely to cover broader viral genotypes and potentially exhibit broad-spectrum antiviral activity across viruses from different families. To assess the antiviral spectrum of Bcl-2 inhibitors, we examined the effect of OLX and ABT-737 on multiplication of two additional mammarenaviruses, the live-attenuated vaccine (LAV) strain, Candid#1, of the New World mammarenavirus Junin (JUNV) (Fig. 6A), and the LASV LAV candidate reassortant ML29, carrying the L segment from the non-pathogenic Mopeia virus and the S segment from LASV (Fig. 6B). For both Candid#1 and ML29 we used their tri-segmented versions expressing the GFP reporter gene (32). In addition, we examined the effect of OLX and ABT-737 on SARS-CoV-2 (Fig. 6C). Both OLX and ABT-737 exhibited a potent dose-dependent antiviral activity against Candid#1, ML29 and SARS-CoV-2.

**Figure 6.**
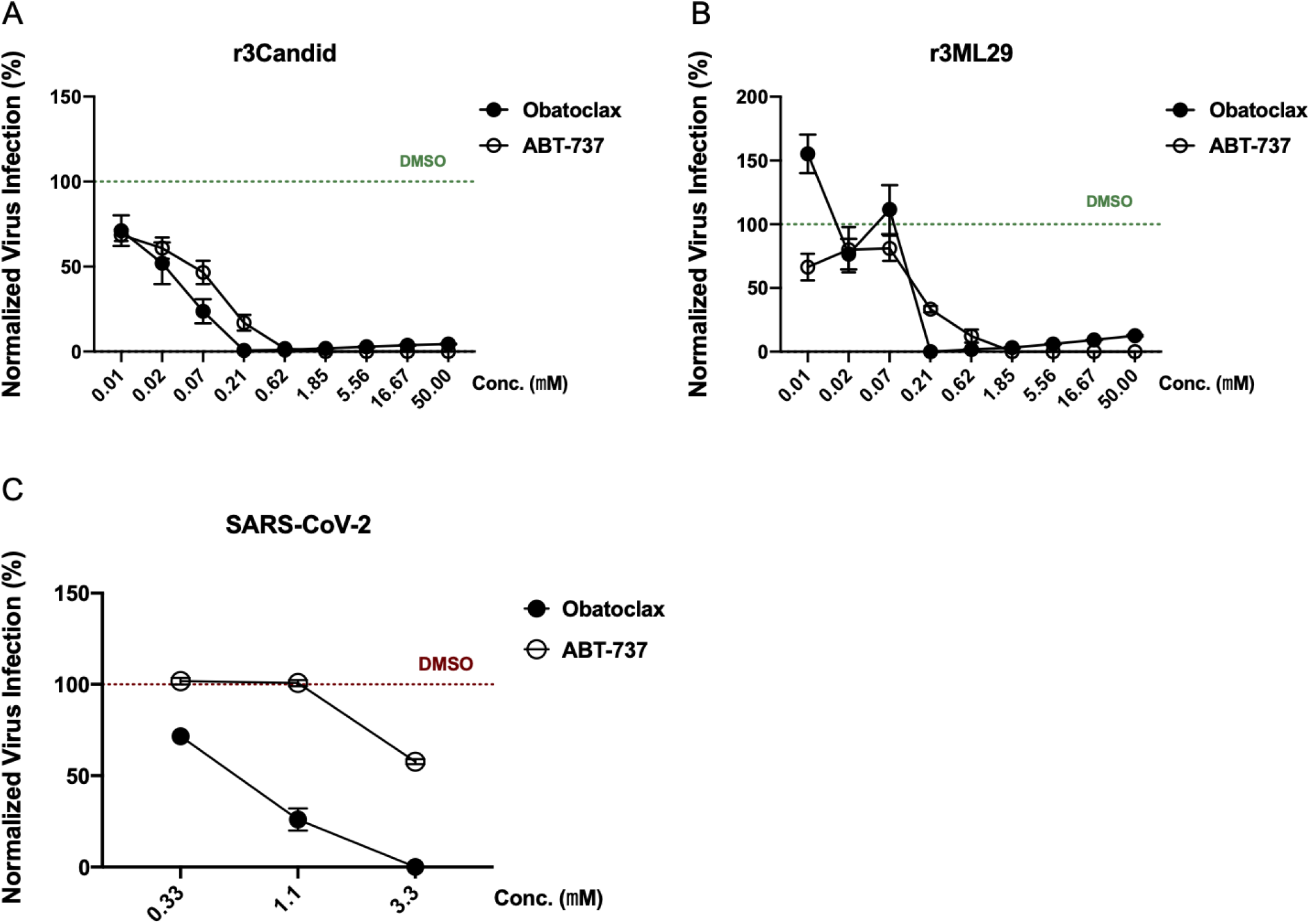
Broad-spectrum antiviral activity of Bcl-2 inhibitors. (A-C) Normalized virus propagation in the presence of OLX or ABT-737. Vero cells were treated with serial dilutions of each compound starting 2 hours prior to infection with r3Can/GFP (A), r3ML29/GFP (B), or SARS-CoV-2 (C). At 72 hpi, GFP expression levels in infected cells (A, B) or titers of infectious SARS-CoV-2 in tissue culture supernatants, as determined by IFFA (C) were measured and normalized to vehicle (DMSO) treated controls.

## DISCUSSION

Host targeting antiviral drugs represent an attractive strategy to combat viral infections, as intra-host virus evolution is unlikely to result in viral variants able to escape from inhibitors that disrupt the activity of host cell factors required for the completion of the virus life cycle. Moreover, related viruses are likely to rely on the same host machinery, thus providing an opportunity for the development of broad-spectrum antiviral therapeutics. Several Bcl-2 targeting small molecules have been identified as potential antiviral drugs against different types of viruses (6-12). However, the mechanisms by which these Bcl-2 inhibitors exerted their antiviral activity are not well understood. In this study, we examined the mechanisms whereby the Bcl-2 inhibitors OLX and ABT-737 exerted their potent anti-LCMV activity. We found that OLX and ABT-737 potent anti-LCMV activity was not associated with their pro-apoptotic properties, but rather their ability of inducing cell cycle arrest at G0/G1 phase.

Small molecules targeting the Bcl-2 family proteins were first developed for cancer treatment, as apoptotic cell death caused by inhibition of anti-apoptotic Bcl-2 proteins contributes to reduce tumor progression (4, 5). We therefore examined whether apoptosis induced by Bcl-2 inhibitors contributed to OLX and ABT-737 anti-LCMV activity. Our finding that OLX and ABT-737 anti-LCMV activity was not affected in the presence of the pan-caspase inhibitor zVAD-FMK or the caspase-3-specific inhibitor zDEVD-FMK (33, 34), indicated that apoptosis did not contribute to the anti-LCMV activity exhibited by the Bcl-2 inhibitors OLX and ABT-737.

In addition to their apoptosis inducing property, Bcl-2 inhibitors have been reported to regulate different cellular processes via mechanisms that are currently poorly understood (29). The role played by Bcl-2 inhibitors in cell cycle progression may also contribute to their tumor-suppressing properties. Accordingly, several anti-cancer compounds that target Bcl-2 have been reported to arrest cell cycle progression, and the corresponding suppression of cell growth shown to play an important role in their anti-tumor activity (13-15, 25). We therefore investigated whether OLX and ABT-737 mediated effects on the regulation of the cell cycle contributed to their anti-LCMV activity. We observed that Bcl-2 inhibitor treatment arrested cells at the G0/G1 phase, which correlated with a significant reduction in expression levels of cell cycle regulators TK1, CCNA2, and CCNB1 (Fig. 3). Consistent with their distinct roles, these cell cycle regulators have been shown to be expressed at distinct phases of cell cycle: TK1 expression starts to increase in the early S phase (35). CCNA is elevated at G2 phase and CCNB is highly expressed at M phase (36). Treatment with OLX and ABT-737 resulted in significant reduction of TK1 expression, consistent with the cells being arrested at G0/G1 phase before entering S phase (Fig. 3B). In contrast, treatment with dinaciclib, known to promote G2/M phase arrest, resulted in highly increased TK1 expression. Our finding that OLX and ABT-737 treatment significantly decreased expression levels of CCNA2 and CCNB1 (Fig. 3B), further supported that expression of G2/M phase regulators was prevented in the presence G0/G1 promoting Bcl-2 inhibitors. Our finding that siRNA-mediated knock-down of cell cycle regulators TK1, CCNA2, and CCNB1 resulted in reduced levels of virus multiplication, suggested that these factors likely play a role in OLX and ABT-737 mediated anti-LCMV activity (Fig. 3C). We found that siRNA-mediated knock-down of Bcl-2 resulted in the strongest reduction on levels of LCMV multiplication, which was associated also with reduced levels of TK1, CCNA2, and CCNB1 (Fig 3D), suggesting that Bcl-2 is an upstream regulator of TK1, CCNA, and CCNB expression.

Several viruses regulate the host cell cycle to create a favorable environment for their replication (16). Thus, influenza A virus (IAV) nonstructural protein 1 (NS1) was found to promote a G0/G1 arrest in infected cells via inhibition of the RhoA-pRb signaling cascade to create favorable conditions for viral replication (37, 38). Likewise, SARS-CoV and the murine CoV mouse hepatitis virus (MHV) have been shown to induce G0/G1 arrest in infected cells to favor their multiplication (39, 40). However, LCMV infection did not affect either cell cycle progression or Bcl-2 inhibitor-mediated G0/G1 arrest (Fig. 3E), and in contrast to IAV and CoVs, cell cycle arrest at G0/G1 generates a cellular environment that contributes to limiting LCMV multiplication. Intriguingly, treatment with ribavirin resulted also in increased numbers of cells at the G0/G1 phase (Fig. 3E), suggesting that cell cycle modulation might be another mechanism, in addition to those previously documented (30, 41) by which ribavirin exerts its broad-spectrum antiviral activity.

We previously showed that OLX did not significantly affect mammarenavirus cell entry or budding, but strongly inhibited steady state levels of mammarenavirus RNA at 48 hpi (17). In the present work we have shown that OLX and ABT-737 did not significantly affect levels of LCMV replication at 2 hpi, but both compounds significantly inhibited viral genomic and anti-genomic RNA synthesis at 4 hpi (Fig. 4). Under our experimental conditions, anti-genomic RNA species consisted of viral cRNA and NP mRNA species. Thus, higher levels of anti-genomic than genomic RNA species observed at 4 hpi likely reflected newly synthesized mRNA. The mechanisms whereby Bcl-2 inhibitor-mediated cell cycle arrest at G0/G1 phase results in reduced levels of RNA synthesis directed by the mammarenavirus polymerase complex remain to be determined.

We selected ABT-737 to examine the anti-LMCV activity of Bcl-2 inhibitors *in vivo*, as its safety and efficacy *in vivo* have been documented for several tumor models (21-24). We observed that at early times after starting ABT-737 treatment, LCMV infected mice showed slightly higher body weight loss compared to vehicle control treated and infected mice (Fig. 5A), suggesting some ABT-737 toxicity. However, after 5 days post-infection, ABT-737 treatment resulted in faster weight recovery compared to the vehicle treated control mice (Fig. 5A). In addition, ABT-737 treatment resulted in reduced serum viral load throughout the entire duration of the observation time with experimental endpoint at 17 days post-infection (Fig. 5B). Toxicity associated with Bcl-2 targeting small molecules can impact their *in vivo* efficacy. Recent studies suggest that implementation of proteolysis-targeting chimera (PROTAC) with Bcl-2 inhibitors could address this issue. PROTACs are modified small molecules in which target molecules are linked to an E3 ligase ligand, promoting ubiquitination of the targets and following degradation through the ubiquitin proteasome system. Notably, converting the Bcl-X_L_ inhibitor, ABT-263, into the Bcl-X_L_ PROTAC form, DT2216, resulted in significantly reduced cytotoxicity but improved anti-tumor activity (42).

OLX and ABT-737 exhibited antiviral activity also against two other mammarenaviruses, ML29, and the most distantly related New World JUNV. In addition, consistent with recent published findings, we found that OLX exerted a strong inhibitory effect on SARS-CoV-2 multiplication (Fig. 6). These results suggest the possibility that Bcl-2 inhibitors actively being explored as anti-cancer therapeutics, could be repositioned as broad-spectrum antivirals.

## MATERIALS AND METHODS

### Cells and viruses

Homo sapiens A549 (ATCC CCL-185), 293T (ATCC CRL-3216), and Grivet (*Chlorocebus aethiops*) Vero E6 (ATCC CRL-1586) cell lines were maintained in Dulbecco’s modified eagle medium (DMEM) (ThermoFisher Scientific, Waltham, MA, USA) containing 10% heat-inactivated fetal bovine serum (FBS), 2 mM of L-glutamine, 100 μg/ml of streptomycin, and 100 U/ml of penicillin. The immunosuppressive strain of LCMV, clone 13 (CL-13) (Reference), recombinant LCMV expressing *Zoanthus* sp. green fluorescent protein (ZsG) fused to nucleoprotein via a P2A ribosomal skipping sequence (rLCMV/ZsG-P2A-NP, referred to as rLCMV/ZsG) (43); the tri-segmented form of the live attenuated vaccine strain Candid#1 of JUNV expressing green fluorescent protein (GFP, r3Can/GFP) (44) have been described previously; the tri-segmented form of reassortant ML29 expressing GFP (r3ML29/GFP) (45) have been described previously. SARS-CoV-2 USA-WA1/2020 (Gen Bank: MN985325.1) was obtained from BEI Resources (NR-52281).

### Treatment of compounds

Cells were plated and incubated at 37 °C and 5% CO_2_ in DMEM containing 2% FBS. After 20 hours, each compound was added at the indicated concentration and for indicated times. Compounds used in vitro experiments were purchased from Selleckchem (Houston, TX, USA); obatoclax; ABT-737; Staurosporine; zVAD-FMK; zDEVD-FMK; dinaciclib. For in vivo test, ABT-737 was purchased from AstaTech Inc (Bristol, PA, USA).

### Cell cytotoxicity assay and half-maximal cytotoxic concentration (CC_50_) determination

Cell viability was assessed using the CellTiter 96 AQueous One Solution Reagent (Promega, Madison). This method determines the number of viable cells based on conversion of formazan product from 3-(4,5-dimethylthazol-2-yl)-5-(3-carboxymethoxyphenyl)-2-(4-sulfophenyl)-2H-tetrazolim by nicotinamide adenine dinucleotide phosphate (NADPH) or nicotinamide adenine dinucleotide phosphate (NADH) generated in living cells. A549 cells were plated on 96-well clear bottom plates (2.0 × 10^4^ cells/well). Serial dilutions (3-fold) of each compound were added to cells, and at 48 hours after drug treatment, CellTiter 96 AQueous One Solution Reagent was added and incubated for 15 min (37°C and 5% CO_2_). Absorbance was measured at 490 nm using an enzyme-linked immunosorbent assay (ELISA) reader (SPECTRA max plus 384, Molecular Devices, Sunnyvale, CA, USA). The resulting optical densities were normalized with dimethylsulfoxide (DMSO) vehicle control group, which was adjusted to 100%. Half-maximal cytotoxic concentrations (CC_50_) were determined using Prism (GraphPad, San Diego, CA, USA).

### Determination of half-maximal effective concentration (EC_50_) and selectivity index (SI)

Cells were plated on 96-well clear-bottom black plates (2.0 × 10^4^ cells/well) and incubated for 20 hours at 37 °C and 5% CO_2_. Cells were pre-treated 2 hours before infection with 3-fold serial dilutions of each compound. Cells were infected (MOI = 0.01) with rLCMV/ZsG-P2A-NP in the presence of compounds. At 48 hours post-infection (hpi), cells were fixed with 4% paraformaldehyde and nuclei were stained with 4′,6-diamidino-2-phenylindole (DAPI), and ZsG expression was determined by fluorescence using a fluorescent plate reader (Synergy H4 Hybrid Multi-Mode Microplate Reader, BioTek, Winooski, VT, USA). Mean relative fluorescence units were normalized with vehicle control group (DMSO), which was adjusted to 100%. EC_50_ were determined using Prism. SIs for hit compounds were determined using the ratio CC_50_/EC_50_.

### Apoptosis determination assays

For Annexin V apoptosis assay, harvested cells were treated with Annexin V-FITC conjugate (BD Pharmingen, Franklin Lakes, NJ, USA) and incubated for 15 minutes at room temperature. After washing stained cells, 7-AAD staining solution (BD Pharmingen) was added and incubate 10 minutes on ice. Cells were detected by flow cytometry (Cytek Aurora, Cytek, Fremont, CA, USA) and data were analyzed using the FlowJo (FlowJo LLC, Ashland, OR, USA). Caspase activity was determined using the Caspase-Glo 3/7 assay system (Promega, Madison, WI, USA) according to the manufacturer’s instruction. The values were normalized with total cellular protein determined with the Pierce BCA Protein Assay Kit (ThermoFisher Scientific).

### Cell cycle studies

To analyze stages of the cell cycle, a BrdU assay kit (APC BrdU flow kit, BD Pharmingen) was used as directed by the manufacturer’s instruction. Briefly, cells were labeled by adding BrdU (final concentration 10μM) and incubated for 1 hour at 37 °C and 5% CO_2_. After harvesting, cells were fixed and permeabilized followed by treatment of DNase to expose incorporated BrdU. Next, cells were stained with APC conjugated anti-BrdU antibody and 7-AAD. Samples were analyzed by flow cytometry (Cytek Aurora) and the FlowJo (FlowJo LLC).

### Western blotting

Whole cell lysates were prepared in RIPA lysis buffer (ThermoFisher Scientific). After sonication for 30 seconds, samples were denatured for 10 minutes at 95 °C. 30 μg of each sample were separated by SDS-PAGE, transferred to the PVDF membrane (Immobilon PVDF membrane, Millipore Sigma, Burlington, MA, USA), and immunoblotted with BCL-2, TK1, CCNA2, CCNB1, and β-actin antibodies (Cell Signaling Technology, Danvers, MA, USA). Bands were visualized with the chemiluminescent substrate (ThermoFisher Scientific).

### Gene knock-down by siRNA

293T cells were transfected with 10 nM of each siRNA using the lipofectamine RNAiMAX transfection reagent (ThermoFisher Scientific), according to the manufacturer’s instruction. Pre-designed ON-TARGET *plu*s siRNA oligoes on the following list were purchased from the Horizon Discovery (Cambridge, UK).

- BCL2 (siRNA ID: J-003307-16)
- TK1 (siRNA ID: J-006787-09)
- CCNA2 (siRNA ID: J-003205-10)
- CCNB1 (siRNA ID: J-003206-09)

### Virus titration

Virus titers were determined by focus-forming assay (FFA) (46). Serial dilutions of samples (10-fold) were used to infect Vero E6 cell monolayers in 96-well plates (2 × 10^4^ cells/well). At 20 h pi, cells were fixed with 4% paraformaldehyde in phosphate-buffered saline. Foci of cells infected with rLCMV/ZsG were determined by epifluorescence of fluorescent reporter gene expression. Foci of cells infected with wild-type LCMV were identified by rat monoclonal antibody VL4 against NP (Bio X Cell, West Lebanon, NH, USA) conjugated to Alexa Fluor 488. SARS-CoV-2 titers were determined by using an immune focus forming assay (IFFA). Vero E6 cells (3×10^4^ cells/well, 96-well plate format, triplicates) were infected with serial viral dilutions (100 µL final volume/96w). At 20 h pi cells were fixed overnight with 10% paraformaldehyde in phosphate-buffered saline. For immunostaining, cells were permeabilized with 0.5% (v/v) Triton X-100 in PBS for 15 min at room temperature and immunostained using the SARS-CoV cross-reactive N protein 1C7C7 monoclonal antibody (1 μg/ml), followed by reaction with a goat anti-mouse conjugated to Alexa Fluor 488.

### RT-qPCR

Cells were infected with rLCMV/ZsG-P2A-NP in the presence of indicated compounds or vehicle control DMSO. Total cellular RNA was isolated using TRI Reagent® (TR 118) (MRC, Cincinnati, OH, USA) according to the manufacturer’s instruction. Total RNA (1 µg) was reverse-transcribed to cDNA using SuperScript™ IV First-Strand Synthesis System (ThermoFisher Scientific). To make strand-specific cDNAs, target primers that are specific to genomic or anti-genomic strand were designed. Target sequences were amplified and quantified by qPCR with primer sets listed in the following section.

### Primers

- (RT reaction)
  - Genomic strand 5’-CAGGGTGCAAGTGGTGTGGTAAGAG -3’
  - Anti-genomic strand 5’-CGAGAACTGCCTTCAAGAGGTCCTC -3’
- (qPCR) LCMV NP
  - Forward 5’-CAGAAATGTTGATGCTGGACTGC-3’
  - Reverse 5’-CAGACCTTGGCTTGCTTTACACAG -3’

### Animal studies

Adult (eight-weeks old) C57BL/6 inbred laboratory mice (Scripps Research breeding colony) were inoculated intravenously (IV) with LCMV Cl-13 (2 × 10^6^ FFU) and treated with ABT-737 (20 mg/kg/day) or vehicle control (30% propylene glycol, 5% Tween 80, and 5% dextrose in water), administered intraperitoneally. Treatment was administered daily from day 1 to 17 post-infection. Mice were monitored daily for development of clinical signs, weight loss, and survival. Sera were collected at day 3, 6, and 17. All animal experiments were done under protocol 09-0137-4 approved by The Scripps Research Institute Institutional Animal Care and Use Committee (IACUC).

## ACKNOWLEDGMENTS

This work was supported by NIH/NIAID grants AI125626 and AI128556. This is the manuscript #30121 from The Scripps Research Institute.

